# Breast cancer incidence as a function of the number of previous mammograms: analysis of the NHS screening programme

**DOI:** 10.1101/238527

**Authors:** Daniel Corcos

## Abstract

The discrepancy between the protective effect of early surgery of breast cancer and the poor benefits of mammography screening programs in the long term can be explained if mammography induces breast cancer at a much higher rate than anticipated. Mammography screening is associated in most countries with a higher incidence of breast cancer, attributed to overdiagnosis. X-ray-induced cancers can be distinguished from overdiagnosed cancers by the fact that their incidence depends on the number of previous mammograms, whereas overdiagnosis solely depends on the last screening mammogram, leading to diagnosis. The unbiased relationship between the number of mammograms and breast cancer incidence was evaluated from the data of the NHS Breast Cancer screening programme in women aged from 50 to 64 years in the United Kingdom. The delay between mammography and increased breast cancer incidence was confirmed from the data of the “Age” trial, a randomized trial of annual screening starting at age 40 in the UK. In women aged 50-64 attending screening at the NHS Breast Cancer programme, *in situ* breast cancer incidence increased linearly from 1993 to 2005 as a function of the number of mammograms. Incidence did not increase anymore after 2005 when the number of mammograms and the delay after screening was stable. Invasive breast cancer incidence increased more specifically in the 60-69 age group. The risk of breast cancer almost doubled after 15 years of screening. Additional cancers began to occur less than 6 years after mammography. These results are evidence that X-ray-induced carcinogenesis, rather than overdiagnosis, is the cause of the increase in breast cancer incidence.

## Introduction

Despite a strong rationale and promising beginnings, the usefulness of mammography screening is now challenged due to poor efficacy in reducing breast cancer (BC) mortality rates [1, 2]. Lack of beneficial effects of mammography screening in the long term contrasts with a better efficacy in short term clinical trials [3–5], where a 31% reduction in mortality from BC has been reported [3]. The lack of change in the incidence of metastatic disease has led to the hypothesis « that BC is a systemic disease by the time it is detectable » [6]. This hypothesis is clearly in contradiction with the protective effect of surgery, which has been known for a century [7, 8], and with the effect of Time To Surgery on cancer mortality [9]. A one-month delay resulted in a 10% added risk of death, with similar added risk for each month delay [9]. As mammography screening results in detection occurring more than one year before clinical cancer [10], it should dramatically reduce cancer mortality. Lack of efficacy of mammography screening in preventing metastatic disease could be explained by the occurrence of a concomitant increase in BC incidence. There is a clear increase in BC incidence worldwide, concomitant to implementation of mammography [11]. However, this increase has been attributed to overdiagnosis [1, 12, 13], i.e. detection by screening of cancers that would not go on to cause death or would spontaneously regress [11]. However, the rate of overdiagnosis is disputed and estimates from random clinical trials suggest than less than five percent of cancers are overdiagnosed [14]. In any case, overdiagnosis does not explain the lack of efficacy of mammography in prevention of metastatic disease [6], nor why screening has not led to a reduction in the incidence of advanced BC [13]. X-rays are known carcinogens [15]. Mammography-induced cancers cannot easily be distinguished from overdiagnosed cancers as both will show an increased incidence in relation to screening. Rates of mammography-induced cancers are expected to be low, an estimate mainly based on extrapolation of data from atomic bomb survivors [16]. However, the validity of the linear no-threshold model is debated [17], and the oncogenicity of low doses still disputed, as DNA repair is not efficient at low doses [18]. Unlike overdiagnosis, x-ray-induced carcinogenesis would explain both the increased incidence of primary BC and the inefficiency of mammography screening.

To evaluate the extent of x-ray-induced carcinogenesis, it is necessary to distinguish x-ray-induced cancers from overdiagnosed cancers. Mammography-induced cancers can be distinguished from the latter by the fact that their incidence will be cumulative, depending on the total number of previous mammograms, whereas overdiagnosis is solely dependent on the screening mammogram leading to diagnosis. To address the distinction between overdiagnosis and x-ray-induced cancers, it is important to use epidemiological data where all the information pertaining to screening is known, and to avoid a selection bias, as women with higher known risk would tend to have higher mammogram numbers.

The NHS BC screening programme began in 1988 on women aged from 50 to 64, and by 1993, all the population of UK within this age group was covered [19, 20]. Each woman had a mammogram once every three years until the age of 64, allowing for tracking of mammogram number. Evaluating the changes in BC incidence as a function of time is complex, because earlier detection as a consequence of screening is a confounding factor. As *in situ* BC are, for the most part, detected by mammography, their reported incidence is a function of the coverage of the age group by screening. Since only a small proportion of them are diagnosed clinically, lead-time bias is not a major concern. The goal of this study was to distinguish raised incidence due to better detection from true increased incidence. After full coverage of the population has been obtained, detection-related changes in the incidence of *in situ* BC incidence should not be observed. In addition, better detection should lead to early detection, i.e. increase in BC found at the prevalent round, but should not lead to major changes in invasive cancer incidence at subsequent screening rounds.

## Materials and Methods

### Information sources

#### The NHS BC screening programme

***In situ* BC incidence**

##### Data collection

Data were collected from documents made publicly available by Cancer Research UK.

The documents used for estimating coverage, uptake (the proportion of screening invitees in a given year for whom a screening test result), and the number of yearly screens of the programme were the following:

Breast Screening Programme, England: 2002-03 [21].

Breast Screening Programme, England: Statistics for 2012-13 [22].

These data covering England (86% of the Great Britain population) were used as an estimate for the UK. Although coverage and uptake changes showed little variation, number of screened women increased during this period, as a consequence of higher invitation rates.

The numbers of women screened each year were the following (millions):

1993: 0.94; 1994: 0.96; 1995: 0.98; 1996: 0.98; 1997: 1.01; 1998: 1.06; 1999: 1.10; 2000: 1.18; 2001: 1.15; 2002: 1.12; 2003: 1.16; 2004: 1.18; 2005: 1.19; 2006: 1.23; 2007: 1.24; 2008: 1.28; 2009: 1.32; 2010: 1.31; 2011: 1.34; 2012: 1.32; 2013: 1.28.

The curves of age-standardized *in situ* BC incidence rates and invasive BC incidence rates can be found on the site of Cancer Research UK [23] and in Supplement Data 1.

Adjustment was done by dividing by the number of women screened in England in the same year.

##### Mammogram counts

The number of mammograms was estimated as follows: With one mammogram every three years, the maximum number of mammograms per woman is five for the 50-64 age group. In 1988, a small percentage of the women born in 1938 and followed until 2002 (when they change the age group) had their first mammogram, and then had one mammogram every three years, the fifth and last in 2000.

For each subsequent generation, the number of mammograms was estimated on the basis of an increment of one mammogram every three years, using a Lexis diagram.

Women of generation 1943, the first with full coverage had their first mammogram in 1993 and the fifth and last in 2005. In 1993, all screened women in the 50-64 age group had one or two mammograms, and from 2000 to 2013, one to five mammograms.

From to 2000 to 2005 the percentage of women with five mammograms should parallel the coverage from 1988 to 1993, with a marked increase in the first three years and a lower increase after.

As the percentage of women with high number of mammograms might have risen slightly from 2003 to 2007 in a way that was difficult to measure, this period was designated as quasi-plateau.

#### Trends in incidence of invasive BC as a function of the age group

Information was found in the Cancer Research UK website [24] and in older sources [25, 26]. The incidence of invasive BC in England and Wales in 1988 (pre-screening period) and 1996 (early screening period) is derived from Figure 5.3, p. 42 of [25]. England and Wales represent 89% of the UK population. It was considered that there should be no major difference relative to the whole UK population.

Incidence in the different age groups in UK in 2005 (late screening period) was found in [26]Epidemiology 3.1.2 Age, Figure 1. Incidence in the different age groups in UK in 2013 is derived from [27] and Supplement Data 2 (BC incidence by age).

**Fig 1.**
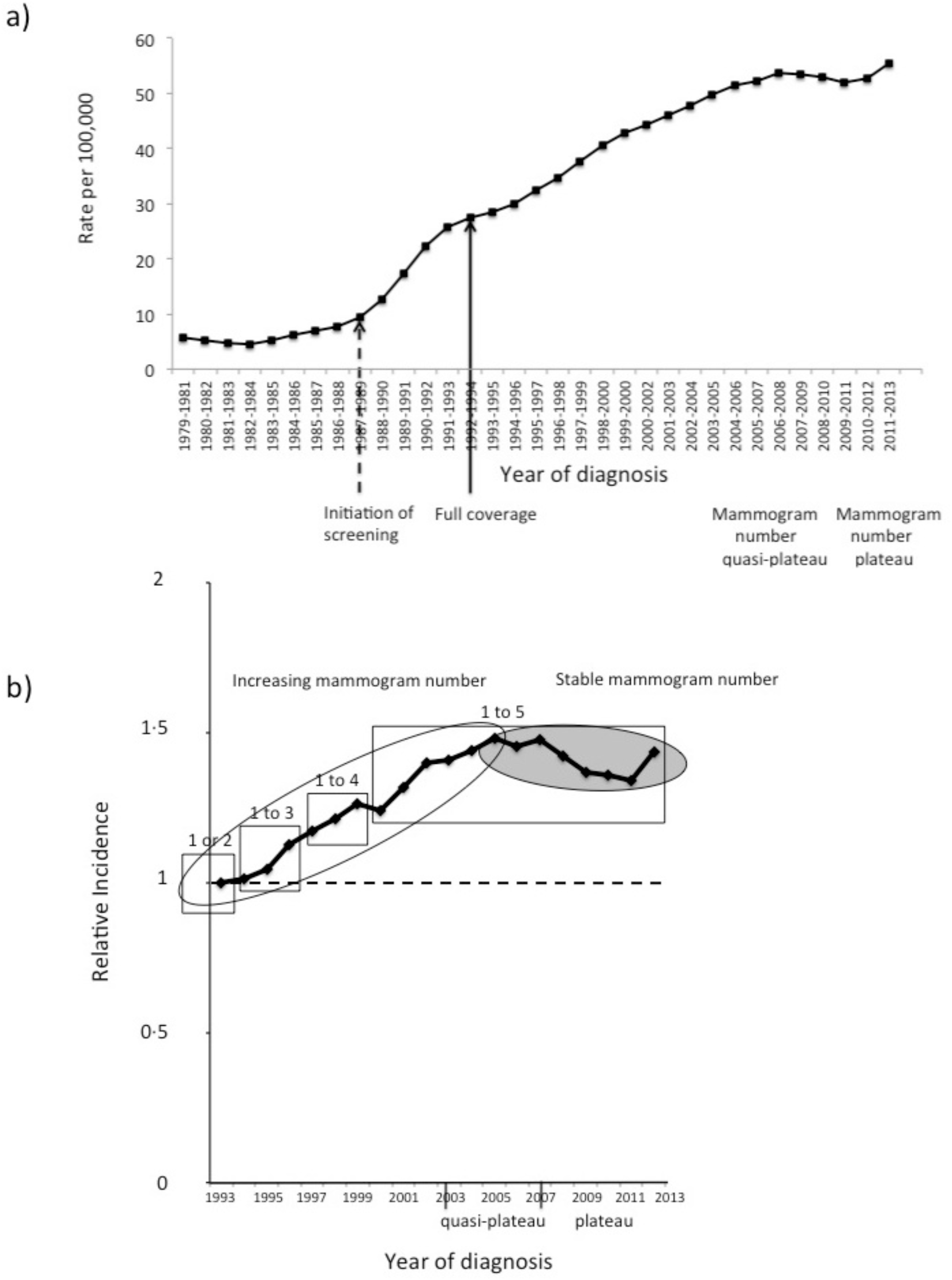
*In Situ* BC Incidence Rates in Great Britain in women aged from 50 to 64 years, 1979-2013, from Cancer Research UK. A) The figure is as reported in the Cancer research UK website for the 50-64 age group. The dotted arrow indicates the beginning of the program in 1988. The solid arrow indicates the year when full coverage of age group 50-64 was attained. Quasi-plateau designates the period where a slight change in the percentage of mammograms of higher rank could occur. B) Incidence rates from figure 1A after adjustment by the screening rate of the population in the corresponding year. Results are expressed relative to the 1993 baseline. Squares indicate the number of mammograms per woman. The open ellipse indicates the period of mammogram number increase for generation 1943, the shaded ellipse the following period.

### The “Age” trial

The « Age » trial [28] was a randomized trial of annual mammographic screening starting at age 40 in the UK. Women were followed for seven years with a mammogram each year [29]. The first screen included 36,348 women. For each following screen, women were included in the study only if they attended all previous screening rounds. 13,998 women attended the last screen. Cancer incidence was monitored at each round of screening, as well as the number of women screened and the mean screening interval.The data were obtained from table 1 of [29]. Adjustment of BC incidence was performed as above, by dividing by the number of women screened at each round, and dividing by the screening interval, which varied from 1.1 to 0.9 year.

**Table 1.**
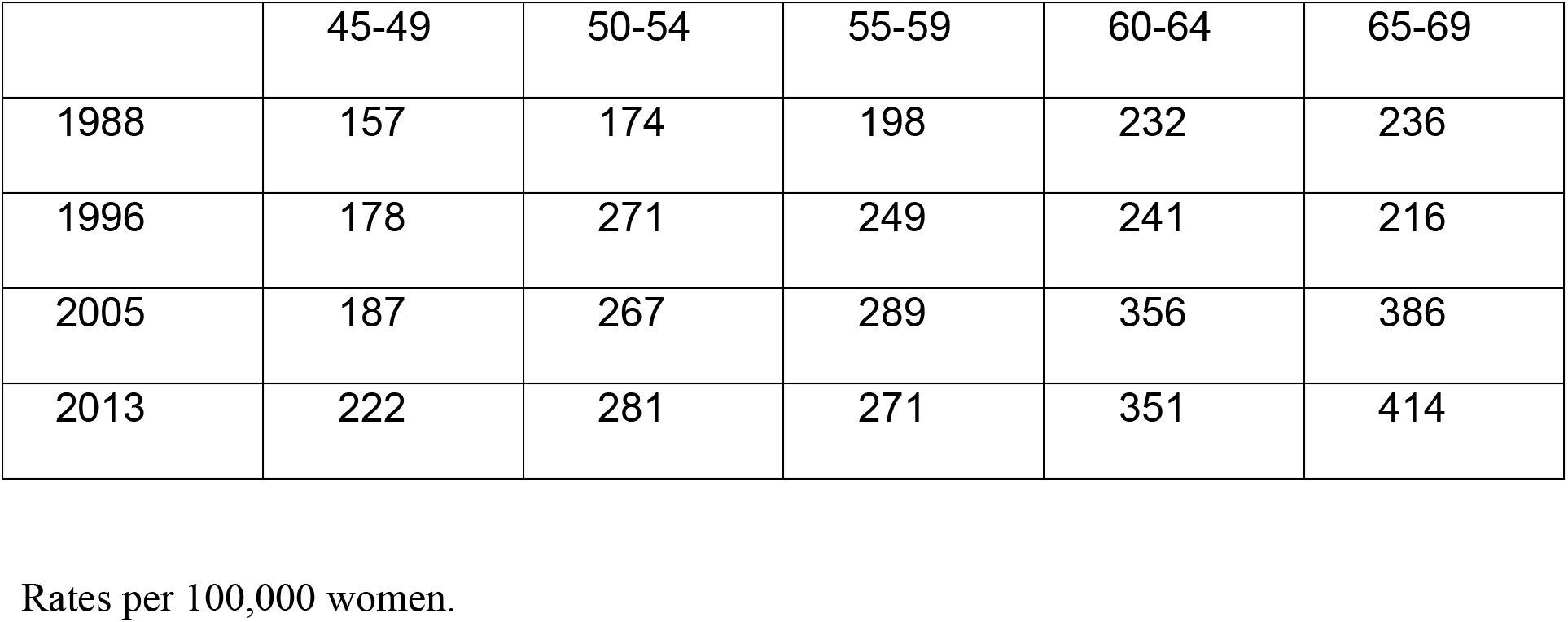
Invasive BC incidence in various age groups at different years.

|  | 45-49 | 50-54 | 55-59 | 60-64 | 65-69 |
| --- | --- | --- | --- | --- | --- |
| 1988 | 157 | 174 | 198 | 232 | 236 |
| 1996 | 178 | 271 | 249 | 241 | 216 |
| 2005 | 187 | 267 | 289 | 356 | 386 |
| 2013 | 222 | 281 | 271 | 351 | 414 |

Rates per 100,000 women.

## Results

In order to avoid the problem of lead-time bias, the study was first focused on *in situ* cancers, since they are detected mainly by screening, as can be seen in figure 1A, where their incidence raises after 1988 at the beginning of screening. In 2005, women of generation 1943, the first generation to get full coverage in 1993, had in average their fifth mammogram. After 2005, the total number of mammograms and the delay after mammograms are almost constant in the 50-64 age group, comprising women ranging in age from 50, having their first mammogram, to women of 64, having had five mammograms. The curve shown in figure 1A indicates that when the total mammogram number and the delay after mammograms remain constant, from 2005 to 2013, the incidence of *in situ* cancers does not vary much with time. The small increase in cases in 2013 is likely to be due to a change in coding rules in England [30].

From 1993 to 2005, the total irradiation of the 50-64 age group increased as a function of time and so did cancer incidence. In this group, the yearly incidence of cancer rose from 27.5 per 100,000 in 1993 to 51.5 per 100,000 in 2005 (figure 1A). Part of this increase could be related to the fact that more women were screened. To eliminate this source of variation, adjustment was performed by dividing the yearly incidence by the number of women screened in England in the same year (figure 1B), from the [21, 22]. With a baseline of 100% corresponding to 1 or 2 mammograms (0 or 1 mammogram before diagnosis) in 1993, screening-adjusted yearly incidence rose by 47.9% in 2005 (figure 1B). Such a dramatic change in incidence was not observed in invasive BC of women outside the screening age [24] during the same period (14·1% for invasive cancer incidence in the 80+ group and 5.9% in the 25-49 age group). No other type of cancer showed such an increased incidence followed by a plateau during the same period in the UK [31]. If the increase of 47·9% in BC incidence that is observed in the mixed population formed by the 50-64 age group is only the consequence of mammograms, then women at the age of 64 having had 12 to 15 years of follow-up and four to five previous mammograms should have almost a double incidence of in situ BC.

Data on age subgroups were lacking regarding *in situ* BC incidence but data were available regarding the trends in invasive BC incidence. For this reason, the impact of previous screening on BC incidence in age subgroups was studied at the stage of invasive BC. The baseline incidence of invasive BC (table 1) was measured from the data in 1988 [25], before the beginning of screening, and the relative changes after screening were evaluated, first in 1996 [25], three years after full coverage, in order to observe the early effect of screening, and at plateau in 2005 [26] and 2013 [27].

As can be seen from figure 2, in 1996, screening was associated with a strong early peak of BC incidence, which can be related to detection of prevalent cancers in the 50-54 age group, and a slight decrease in invasive BC in the 65-69 age group, as expected from the aim of « beating cancer sooner ». However at plateau, from 2005 to 2013, while the early peak (the 50-54 age group) was of the same magnitude, indicating no major change in baseline, relative incidence of BC in the 60-64 and the 65-69 groups was equal or higher than that in the early peak, indicating that not only the aim of «beating cancer sooner» was not reached, but that an excess risk of late cancer was present. The change in the 60-64 age group is particularly meaningful since this group was covered by screening from 1993, and therefore a prevalent screening round is unlikely to explain this increase. The magnitude of the increase is in agreement with the estimates for *in situ* BC.

**Fig 2.**
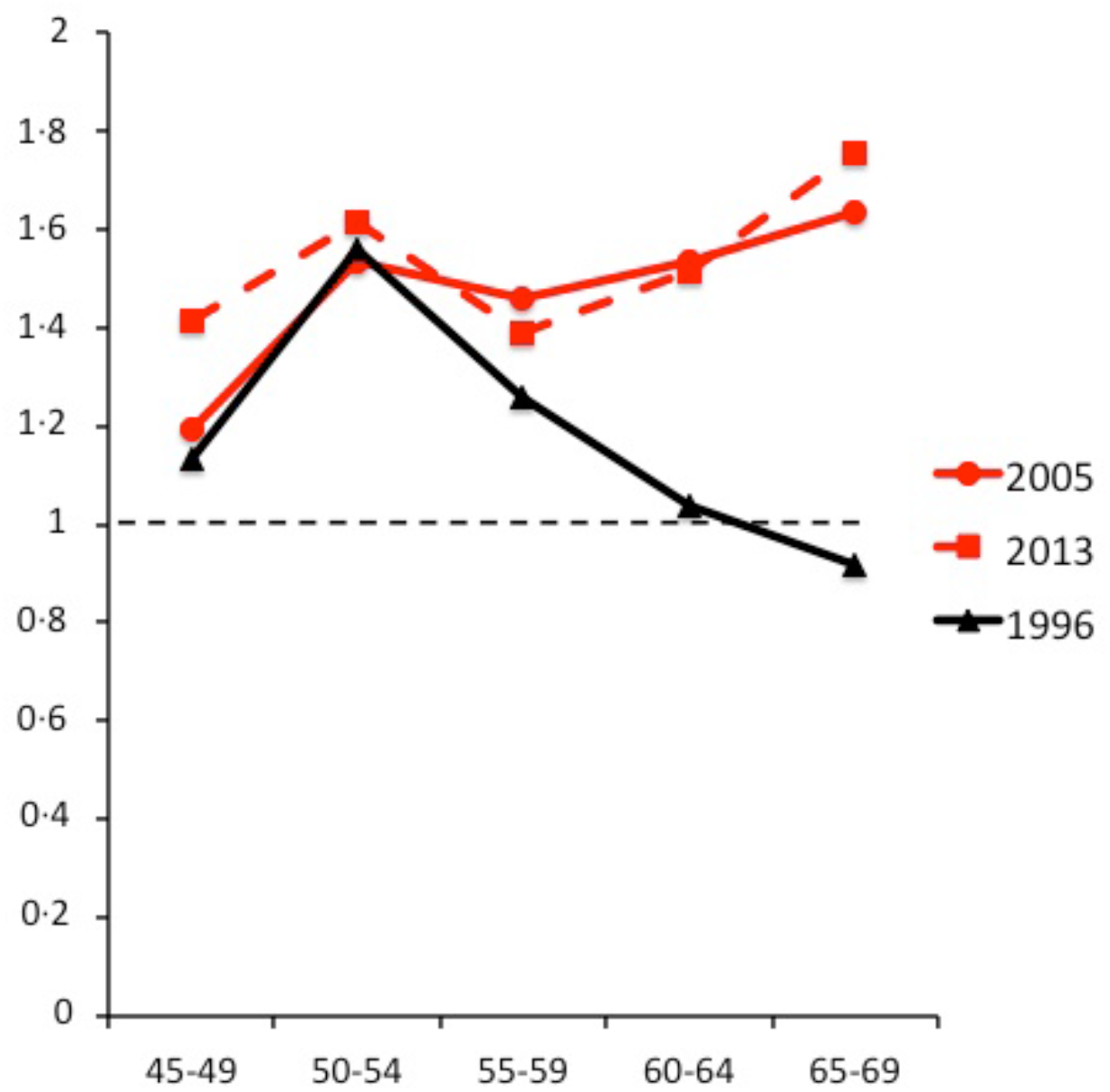
Invasive BC incidence relative to pre-screening levels in various age groups at different years. Yearly incidence in the age groups at the indicated years was divided by yearly incidence in 1988, before screening (from Table 1). The baseline is indicated by the black dotted line. Early effects of screening (black curve) can be compared to late effects (red curves).

The extent of change in incidence of invasive BC can be assessed in the 65-69 age group: after a period of incidence stability between 1993 and 1996, this group showed a strong increase in BC incidence, mainly from 2001 to 2007 (figure 3), consistent with exposure to mammography in previous years. The increase in incidence precedes the nationwide implementation of screening in the 65-70 age group and a plateau 64% above the basal level is observed between 2006 and 2013. As the incidence begins to increase before the first screening year of this age group and as the high level is persistent, this excludes the possibility that it is due to lead-time bias (which is also unlikely because these women were already screened from their fifties). From this curve, it can be concluded that birth cohorts born before 1927, which could not have had the full effect of screening, had a much lower incidence of BC than cohorts born after 1941, which have had the maximum effect of screening. In two other age groups, which were not affected by national screening, BC incidence was relatively stable (figure 3).

**Fig 3.**
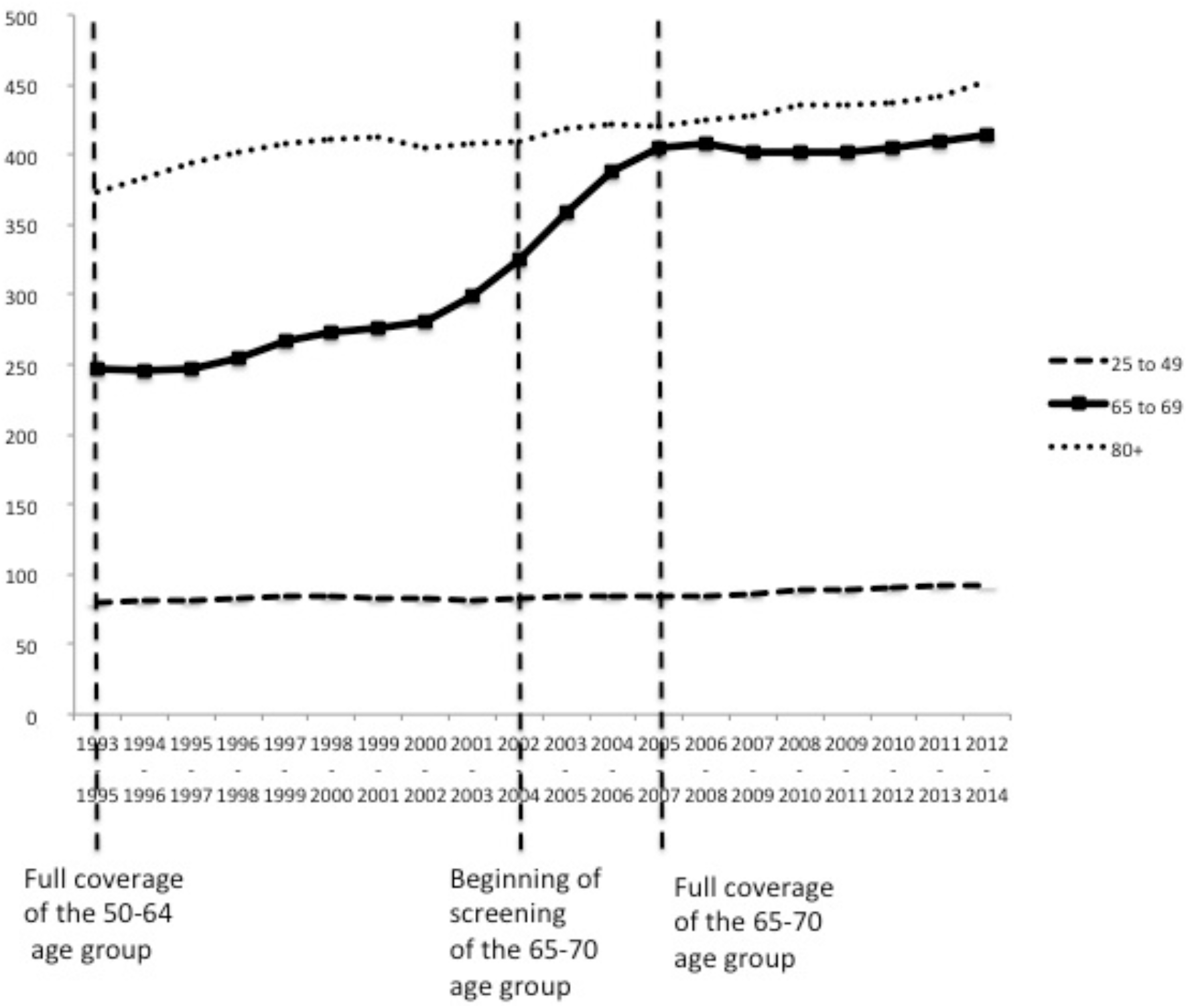
Invasive BC incidence rates per 100,000 women in the United Kingdom for different age groups, from Cancer Research UK. Events related to screening implementation are indicated by vertical dotted lines. The solid curve represents BC incidence in the 65-69 age group, affected by previous screening. The dotted curves represent the 20-49 age group and the 80+ age group, unaffected by screening.

The latency of cancer occurrence after mammography appears to be short, as a rise in *in situ* BC incidence can be detected in 1994, 6 years after the beginning of screening implementation. The short latency after irradiation opened the possibility to make an independent assessment of the effect of mammograms in short term clinical trials. Women in the intervention arm of the « Age » trial [28] in the UK had annual mammographic screening starting at age 40 and were followed for eight years. As expected, since the delay after irradiation was too short for allowing the growth of a tumor large enough to be detected by mammography, *in situ* BC incidence was stable during the first screening rounds, then rose strongly at the 6th mammogram corresponding to the fifth year of screening (figure 4A, black bars). Similarly, the increase in invasive BC incidence (figure 4A, open bars) was delayed after the beginning of screening. Unlike the NHS BC screening programme, this analysis could have been biased, as women with a family history of BC would be more likely to continue in the study, which would translate in a higher BC incidence in the last rounds. However, drop-out was more important during the first five rounds than during the last three ones (figure 4B). It is therefore unlikely that the observed results are due to self-selection. It is also unlikely that an age-related change could account for all the observed results, as it would mean a doubling in BC incidence between the age of 45 and 47 and almost no change before. Instead, these results suggest a lag before an x-ray-induced cancer is large enough to be detected. Nevertheless, the age-related changes should be taken into account to evaluate more precisely the level of early induction of cancer, as the age-specific incidence in the 45-49 age group is about 1.5 greater than that in the 40-44 age group [25, 26].

**Fig 4.**
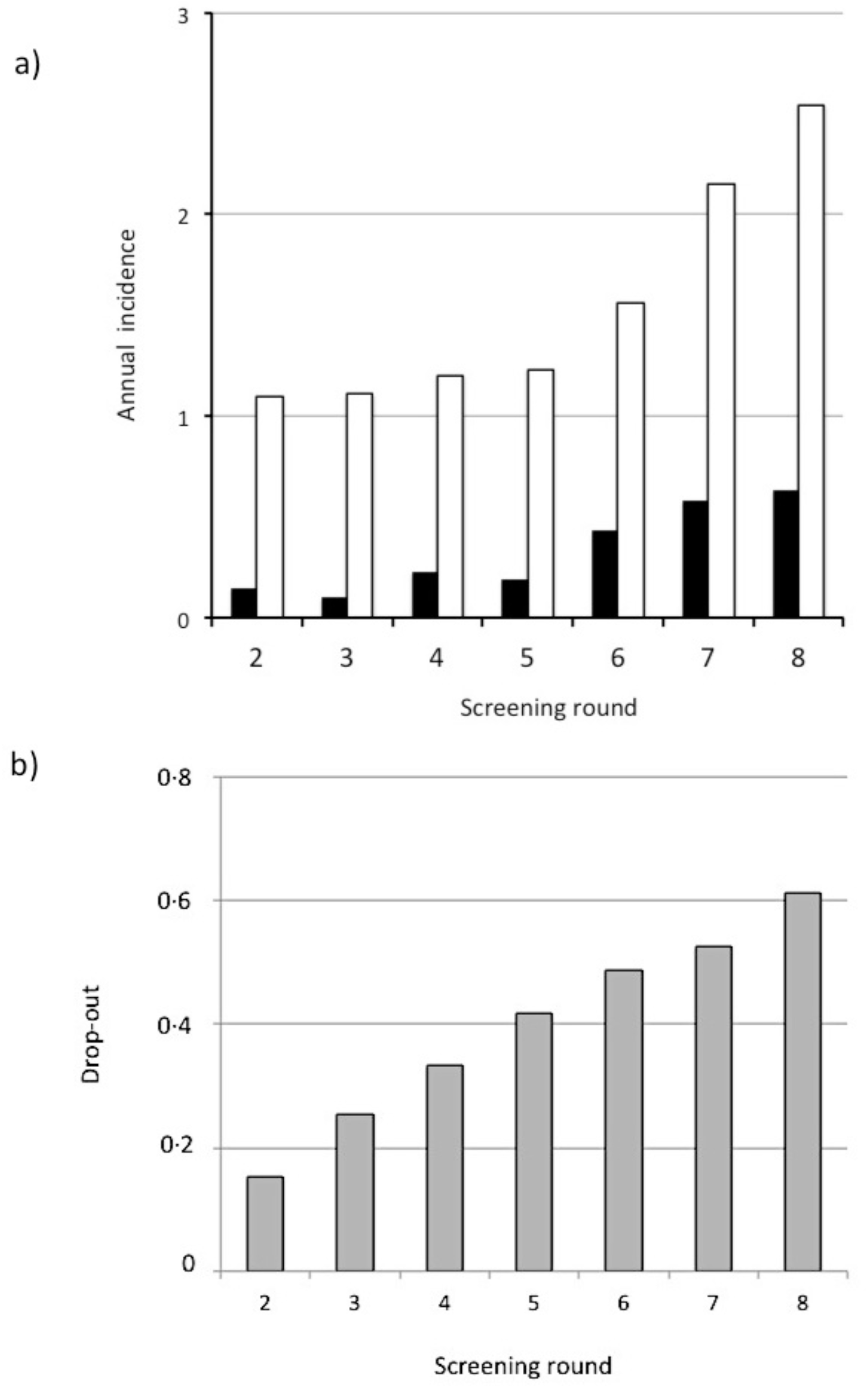
BC incidence as a function of screening round, from the “Age” trial. A) Results are expressed as annual incidence per thousand women screened, from the results of Gunsoy et al. [29], after adjustment by the screening interval. Mean screening interval is one year. First round of screening was excluded as it corresponds to prevalent cancers. Total number of *in situ* cancers: 44. Total number of invasive cancers: 226. Black bars: *in situ* cancers. Open bars: invasive cancers. B) Drop-out as a function of the screening round. Results are presented as the ratio of the number of women having left the trial to the number of women at first round.

## Discussion

The aim of this study was to test the hypothesis that the raised incidence observed worldwide concomitant to mammography screening would be due to the accumulation of low-dose x-ray radiations by the breasts. For this, it was necessary to eliminate artificial increase due to better detection. This study indicates that *in situ* BC incidence was correlated to the number of previous mammograms in women aged 50-64 years who underwent mammography screening from 1993 to 2013. One may consider the unlikely possibility that there was a trend in mammogram sensitivity that was incremental during 1993 and 2005 and then reached a plateau. However, in the US, where screening had been implemented earlier, there was no increase in *in situ* cancer incidence after 1998 that would suggest a better detection by mammograms [32]. One could argue that the increase of BC incidence in UK is related to a better detection of cancer due to technical improvement with time in this country, or to a fortuitous concomitant increase of BC incidence at exactly the same time. This seems highly unlikely, as BC incidence did not change between 1996 and 2005 for the 50-54 age group, and changed dramatically for the 60-69 age group, who had experienced mammography screening (figure 2). An unbiased relationship between previous mammogram number and cancer incidence suggests that x-ray-induced carcinogenesis and not overdiagnosis is involved in the increase of BC incidence after implementation of mammography screening. The age stratification of the women in the NHS BC programme years avoids age related biases. Also, one would expect overdiagnosis occurring predominantly on prevalent cancers (the 50-54 age group), not on incident cancers (other age groups). Therefore, the specific increase observed between 1996 and later years in women older than 60 years cannot be related to overdiagnosis. As a long-lasting excess of BC after mammography screening is observed in all countries after implementation of mammography screening [11], it is very unlikely that important life style changes that may affect BC incidence would be temporally associated with screening in each of these countries. One would also have to explain why life style changes would have so few effects in UK women who did not have the time to develop x-ray-induced cancers (the raised incidence between 1988 and 1996 in the 50-54 age group is accounted for by detection of prevalent cancers). In addition, the risks related to life style, like obesity and age at first live birth, are not large enough [33] to account for the observations in the 60-69 group. A similar increase in BC incidence in the 60-69 age group is observed in other countries late after implementation of mammography screening (Corcos, in preparation). Finally, it would be possible that a common cause, like familial risk, would affect both previous mammogram number and cancer incidence in a sample population, but not in a nationwide screening program.

Therefore the relationship between number of previous mammograms and BC incidence suggests direct causation, a conclusion which is supported by the fact that the number of excess cancers in screened population increases with time (Corcos, submitted). Ethical issues preclude more detailed studies addressing the risk related to one single mammogram. However, extension of the NHS Breast Screening programme to women aged 47 to 49 after randomization [34] should provide valuable information.

Although the effect of mammography screening on cancer incidence could have been predicted from the discrepancy between the poor efficiency of population mammography screening on metastasis rate [6] and its expected outcome, particularly from short term clinical trials [3, 4], it was not anticipated from current risk models [16]. Although the linear no-threshold model is questioned, most authors conclude to the safety of low doses [35, 36], with the notable exception of Gofman, who claimed that medical x-rays were responsible for 75 percent of breast cancers in the USA [37]. There is clear evidence for strong mutagenic effects of ultra-low doses at high rate exposure [38], which might be relevant to mammography screening.

The 5 to 7 years latency observed in both studies is at variance with the popular view of a long latency of x-ray-induced cancers. However, if it is true that x-rays are more carcinogenic at the time of puberty and that in this case, cancers arise with a long delay, irradiated women aged over 50 at the time of bombing showed increased occurrence of cancers that was not delayed as compared to non-exposed women [39]. An excess of cumulative cancer incidence of 22% has been found as early as 6 years after the first round of mammography in screened women as compared to a control group, but this difference has been attributed to spontaneous regression of BC [11], in contradiction with the clinical observations before the surgery era [7, 8]. Therefore it is likely that the excess observed was related to mammography-induced cancers.

DNA damage, and particularly the formation of DNA double strand breaks (DSB), is the main factor in the carcinogenic action of X-rays [15, 40]. Many BC predisposition genes encode proteins involved in DNA DSB repair pathways [41]. Some studies have suggested a positive association between radiation exposure and BC risk in BRCA1/2 mutation carriers [42, 43], while others did not find significant effects [44, 45]. However, the latter studies were heavily biased (Corcos, in preparation). It would be interesting to know whether the contribution of DNA-damage repair genes to global BC burden has increased following mammography-screening implementation.

In the context of the overdiagnosis paradigm, the absence of an effect of mammography screening on the rate of metastatic cancer does not encourage use of other methods for screening. Alternatively, if we adhere to the principle that early diagnosis and prompt treatment save lives, which can be consistent with the present findings, screening must be performed. From a public health point of view, it might be a concern that the last mammogram of a screening is not followed by detection of the cancer it may have caused. The best way to address the problems created by mammography is to replace it by a safer method for first-line screening. With some exceptions, Magnetic Resonance Imaging (MRI) is not as useful as mammography for detecting BC [46]. However, as it is unlikely to be carcinogenic, replacement of mammography screening by MRI screening should result in a strong decrease in BC mortality rates.

## Acknowledgments

I thank Vasco Rodrigues, Vincent Favaudon, and Michael Osborn for comments on the manuscript.

## Supporting Information

**Supplemental data 1: Incidence of *in situ* Breast Cancer (1979-2013)**

From Cancer Research UK (2016)

**Supplemental data 2: Invasive Breast Cancer incidence by age (2012-14)**

From Cancer Research UK (2016)

